# Causal evidence for mnemonic metacognition in human precuneus

**DOI:** 10.1101/280750

**Authors:** Qun Ye, Futing Zou, Hakwan Lau, Yi Hu, Sze Chai Kwok

**Affiliations:** Shanghai Key Laboratory of Brain Functional Genomics, Key Laboratory of Brain Functional Genomics (Ministry of Education),School of Psychology and Cognitive Science, East China Normal University, Shanghai200062, China; Shanghai Key Laboratory of Magnetic Resonance, East China Normal University, Shanghai 200062, China; NYU-ECNU Institute of Brain and Cognitive Science at NYU Shanghai, Shanghai 200062, China; Department of Psychology, University of California-Los Angeles, Los Angeles, California, 90095, United States; Brain Research Institute, University of California-Los Angeles, Los Angeles, California, 90095, United States; Department of Psychology, University of Hong Kong, Hong Kong

## Abstract

Metacognition is the capacity to introspectively monitor and control one’s own cognitive processes. Previous anatomical and functional neuroimaging findings implicated the important role of precuneus in metacognition processing, especially during mnemonic tasks. However, the issue of whether this medial parietal cortex is a domain-specific region that supports mnemonic metacognition remains controversial. Here, we focally disrupted this parietal area with repetitive transcranial magnetic stimulation in healthy participants of both sexes, seeking to ascertain its functional necessity for metacognition for memory versus perceptual decisions. Perturbing the precuneal activity impaired the metacognitive efficiency selectively in the memory judgment of temporal-order, but not in perceptual discrimination. Moreover, the correlation in individuals’ metacognitive efficiency between the domains disappeared when the precuneus was perturbed. Together with the previous finding that lesion to the anterior prefrontal cortex impairs perceptual but not mnemonic metacognition, we double dissociated the macro-anatomical underpinnings for the two kinds of metacognitive capacity in an interconnected network of brain regions.

**SIGNIFICANCE STATEMENT:** Theories on the neural basis of metacognition have thus far largely centered on the role of prefrontal cortex. Here we refined the theoretical framework through characterizing a unique precuneal involvement in mnemonic metacognition with a noninvasive but inferentially powerful method: transcranial magnetic stimulation. By quantifying meta-cognitive efficiency across two distinct domains (memory vs. perception) that are matched for stimulus characteristics, we reveal an instrumental – and highly selective – role of the precuneus in mnemonic metacognition. These causal evidence corroborate ample clinical reports that parietal lobe lesions often produce inaccurate self-reports of confidence in memory recollection and establish that the precuneus as a nexus for the introspective ability to evaluate the success of memory judgment in humans.

## INTRODUCTION

Metacognition is the ability to introspectively monitor and control one’s own cognitive processes, which is important to guide adaptive behavior, social interaction and mental health (Flavell, 1979; Frith, 2012; Nelson, 1990; Teasdale et al., 2002). Metacognitive capacity has been mostly assessed by self-reporting of level of confidence in one’s own decisions that correlate with objective performance. The initial task is often called “type 1 task” and the ensuing confidence judgment task is called “type 2 task” (Galvin, Podd, Drga, & Whitmore, 2003). A widely used approach to estimate the metacognitive efficiency without having it confounded by the primary task performance and response bias is to calculate the comparison between the type 1 sensitivity (d’) and the type 2 sensitivity (meta-d’). This approach can quantify meta-ability under the signal detection theory (SDT) framework (Maniscalco & Lau, 2012) or by a recently developed hierarchical Bayesian estimation method (Fleming, 2017).

Despite a large amount of recent research showing the neural architecture of metacognition in various cognitive domains, like visual perception and memory (Baird, Smallwood, Gorgolewski, & Margulies, 2013; Fleming, Ryu, Golfinos, & Blackmon, 2014; Fleming, Weil, Nagy, Dolan, & Rees, 2010; McCurdy et al., 2013; Rahnev, Nee, Riddle, Larson, & D’Esposito, 2016; Yokoyama et al., 2010), the underlying mechanisms of metacognition are incompletely understood. A central question is whether the human metacognition depends on some domain-general neural structures, or is it supported by domain-specific components? While it has been reported that metacognitive behavioral indices are correlated across the memory versus perception domains (Faivre, Filevich, Solovey, Kuhn, & Blanke, 2016; McCurdy et al., 2013), their functional neural correlates might be largely independent (Baird et al., 2013).

In contrast to the established role of the anterior prefrontal cortex in perceptual metacognition (Fleming et al., 2014), compelling evidence converge to reveal an important role of the precuneus in memory metacognition (Fleck, Daselaar, Dobbins, & Cabeza, 2006; Fleming et al., 2010; McCurdy et al., 2013). Functionally, it has been shown that the task-related activity in the precuneus was greater during memory task compared to during perceptual decision (Morales et al., 2018). Anatomically, the structural variation in the precuneal region was correlated more robustly with memory metacognitive efficiency than with visual perceptual metacognitive efficiency, ascribing a critical role of the precuneus in meta-memory (McCurdy et al., 2013).

The extant evidence for the function of precuneus in metacognition has been correlational. Here we used a disruptive technique that can non-invasively establish the causal role of the precuneus in metacognition across the memory and perceptual domains. We applied transcranial magnetic stimulation (TMS) over either on the precuneus or a control site before the type 1 tasks to perturb the neural activity so as to ascertain whether the precuneus might be causally involved in metacognition in either or both domains. In both tasks, on each trial the participants were required to make a two-alternative forced choice judgment between a pair of still frames, followed by a confidence rating of their choice decision for that trial; the only difference between the two tasks was the task-demands. In the memory task, the participants were asked to identify the image that was presented earlier in a video gameplay that they had encoded 24 hours earlier; in the visual perceptual task, the same group of participants were required to discriminate the difference in resolution between the two images. We kept the individual sets of pair-images identical in both tasks per participant. To anticipate, we expected a Task × TMS interaction, which shall arise from a more pronounced deficit in the meta-memory efficiency following TMS to the precuneus than in the meta-perceptual efficiency.

## MATERIALS AND METHODS

### Participants

18 adults (7 female, age 19-24 years) from the student community of the East China Normal University participated in this study. Each of them participated in both tasks, giving us a within-subjects comparison. All participants had normal or corrected-to-normal vision, no reported history of neurological disease, no other contraindications for MRI or TMS, and all gave written informed consent. They were compensated financially for their participation. No subject withdrew due to complications from the TMS procedures, and no negative treatment responses were observed. The study was approved by University Committee on Human Research Protection of East China Normal University (UCHRP-ECNU).

### Overview of study

The memory task and perceptual task were separated into two experimental sessions. Immediately before performing the main task, the participants received 20 min of repetitive TMS that targeted at one of the two cortical sites (Within-subjects: TMS-vertex vs. TMS-precuneus) in a counter-balanced manner (Experimental session 1 and session 2 in Figure 1A). The session order, the numbers of trials, the dimension, position and sequence of stimulus presented, the response time allowed for the type 1 task judgment and the inter-trial intervals (ITIs) were all identical in both tasks. High-resolution structural scans were acquired for each participant to guide the TMS procedure.

**Figure 1:**
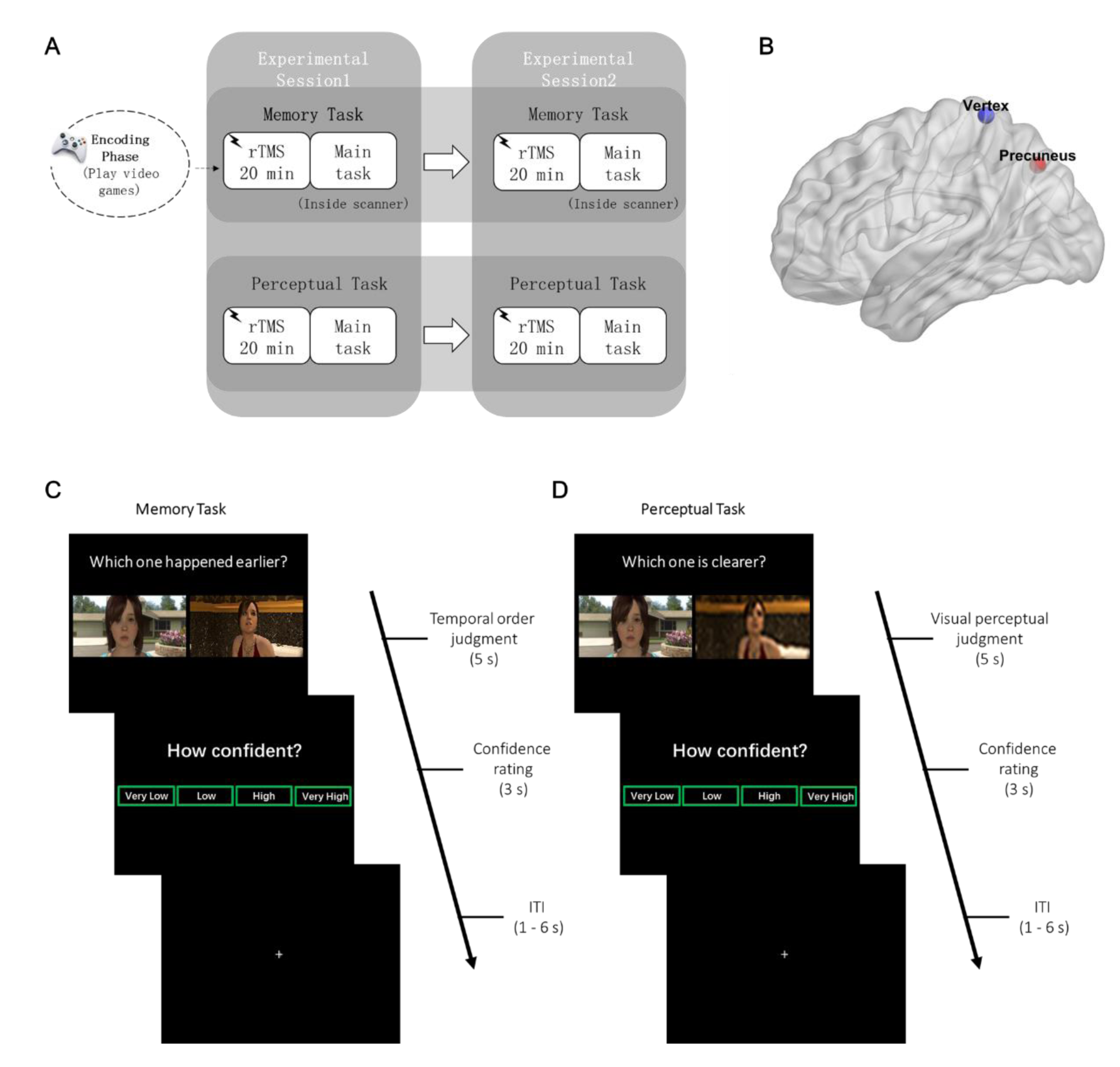
Study design. **(A)** Experimental overview. In experimental sessions 1 and 2 of both tasks, participants received 20 min of rTMS to either one of two cortical sites before performing the main task. The stimulation sites (within-subjects design: TMS- precuneus vs. TMS-vertex) and choices of video game chapters were counterbalanced within subjects across task. **(B)** Location of precuneus (target site) is depicted in red and vertex (control site) in blue. The target site for precuneus stimulation (MNI x y z = 6, −70, 44) was based on (Kwok, Shallice, & Macaluso, 2012). **(C)** In memory task, the participants performed a temporal order judgment task, by choosing the image that happened earlier in the video game. **(D)** In perceptual task, participants identified which frame out of the two was clearer (or blurrier). After the type 1 tasks, participants rated their confidence level on a 4-point scale.

### Experimental Design and Statistical Analysis

#### Tasks and procedure

Each participant completed 480 trials in total in each of the two tasks (2 sessions × 4 blocks × 60 trials per block).

The memory task required participants to choose the image that happened earlier (temporal order judgment, TOJ) in the video game they had played one day before. The retrieval task was administrated inside an MRI scanner, where visual stimuli were presented using E-prime software (Psychology Software Tools, Inc., Pittsburgh, PA), as back-projected via a mirror system to the participant. Each trial was presented for 5 s during which participants performed the TOJ. They were then allowed 3 s to report their confidence level following the memory judgment. Participants performed the TOJ task using their index and middle fingers of one of their hands via an MRI compatible five-button response keyboard (Sinorad, Shenzhen, China). The participants reported their confidence level (“Very Low”, “Low”, “High”, or “Very High”) regarding their own judgment of the correctness of TOJ with four fingers of the other hand. The left/right hand response contingency was counterbalanced across participants. The participants were encouraged to report their confidence level in a relative way and make use of the whole confidence scale. Following these judgments, a fixation cross with a variable duration (1 – 6 s) was presented (Figure 1C).

The same sets of paired-images were used in the perceptual task, in which the participants were required to choose either the clearer (or blurrier, counter-balanced across participants) image among a pair of images on each trial. The participants made an image-resolution comparison judgment and then a confidence rating of their type 1 task decision (Figure 1D) with a 17-inch CRT monitor in a dimly illuminated room. There was a practice block before each session for the participant to get familiar with the task demands.

#### Quantification of metacognitive efficiency

Memory and perceptual performance were quantified using the percentage of correct judgments and the d’ of type 1 signal detection theory (Green and Swets, 1966; Macmillan and Creelman, 2004). We evaluated the metacognitive ability of both tasks by meta-d’. Meta-d’ quantifies metacognitive sensitivity (the ability to discriminate between correct and incorrect judgments) in a signal detection theory (SDT) framework. Meta-d’ was widely used as a measure of metacognitive capacity because it is expressed in the same scale as d’, so the type 2 sensitivity (meta-d’) could be compared with the type 1 sensitivity (d’) directly (Fleming & Lau, 2014; Maniscalco & Lau, 2012). If meta-d’ equals to d’, it means that the metacognitive sensitivity is ideal. Here, we calculated the M-diff (meta-d’ minus d’) for estimating the metacognitive efficiency (the level of metacognition given a particular level of performance or signal processing capacity). The toolbox on MATLAB for the SDT-based meta-d’ estimation was available at http://www.columbia.edu/∼bsm2105/type2sdt/.Moreover,we computedthe metacognitive efficiency using a hierarchical Bayesian estimation method (https://github.com/smfleming/HMeta-d), which can avoid edge-correction confounds and enhance statistical power (Fleming & Daw, 2017). The 4-point confidence ratings were collapsed into two categories (high and low) for all analysis.

Additionally, to ensure our results were not due to any idiosyncratic violation of the assumptions of SDT, we calculated the phi coefficient index, which represents each subject’s correlation between their discrimination accuracy and confidence ratings (Kornell, Son, & Terrace, 2007). The phi coefficient was calculated according to the following formula using the number of trials classified in each case [n(case)]:

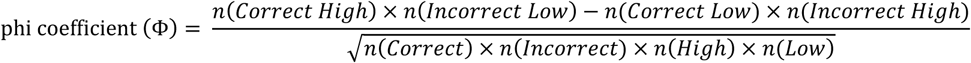

Data were processed with in-house software on MATLAB and statistical inference was made using Rstudio.

### Stimuli

The stimuli were extracted from an action-adventure video game (Beyond: Two Souls), which was created by the French game developer Quantic Dream and played in the PlayStation 4 video game console developed by Sony Computer Entertainment. The Participants played 14 chapters in total across two sessions: 7 in experimental session 1 and then another 7 in session 2. These subject-specific video were recorded and were used for extraction of still images for the tasks.

For the memory task, we selected static images from the subject-specific recorded videos which the participants had played the day before. Each second in the video consisted of 29.97 static images (frames). In each game-playing session, 240 pairs of images were extracted from the seven chapters and were paired up for the task based on the following criteria: (1) the two images had to be extracted from either the same chapters or adjacent chapters (Within-vs. Across-chapter condition); (2) the temporal distance (TD) between the two images were matched between Within- and Across-chapter condition; (3) in order to maximize the range of TD, we first selected the second longest chapter of the video and determined the longest TD according to a power function (power = 1.5), at the same time ensuring the shortest TD to be longer than 30 frames. We generated 60 progressive levels of TD among these pairs.

For the perceptual task, the same sets of subject-specific stimuli from the memory task were used. On each trial, the resolution of one of the images was reduced using Python Imaging Library through resizing the image to change the pixel dimension. For instance, setting an image to three-tenths of the original size changed the pixel dimension to three-tenths, then the image was resized to its primary size so that the pixels per inch (PPI) decreased proportionately. The higher the PPI, the smaller the difference between the image resolution of the resized one and the original was, which also meant this pair would be harder to discriminate than another pair with a lower PPI value. Based on participants performance in the memory task, we pre-determined five difficulty levels for the perceptual task (n = 1∼5, 1 is the hardest). The image resolution was adjusted online using an n-down/1-up adaptive staircase procedure, aiming to equate individual performance with his or her performance in the memory task.

### Anatomical MRI images

A 3-Tesla Siemens Trio magnetic resonance imaging scanner (Siemens Medical Solutions, Erlangen, Germany) was used to acquire the high-resolution T1-weighted images for each participant (192 sagittal slices, TR = 2530 ms, TE = 2.34 ms, TI = 1100 ms, flip angle = 7°, FOV = 256 × 256 mm, 0.9 mm thickness, voxel size = 1 × 1 × 1 mm) to stereotaxically guide the transcranial stimulation.

### Repetitive transcranial magnetic stimulation (rTMS): procedure, protocol and sites

TMS is a form of noninvasive cortical stimulation method that can modulate cognitive functions. Previous studies have demonstrated that repetitive stimulation with TMS over the precuneus (Kraft et al., 2015) or lateral parietal cortices (Nilakantan, Bridge, Gagnon, VanHaerents, & Voss, 2017; Wang et al., 2014) produce robust effects on memory related ability, showing the efficacy of rTMS targeted at relatively deep regions. The present study adopted the identical stimulation magnitude and protocols used in our previous study (Ye, Hu, Ku, Appiah, & Kwok, 2018).

The rTMS was applied using a Magstim Rapid magnetic stimulator connected to a 70mm double air film coil (Magstim Company). The structural T1-weighted magnetic resonance images were obtained for each subject and used in the Brainsight2.0, a computerized frameless stereotaxic system (Rogue Research), to localize the target brain regions. Target stimulation regions for rTMS were selected in the system by transformation of the Montreal Neurological Institute (MNI) stereotaxic coordinates to participant’s normalized brain. The sites stimulated were located in the precuneus at the MNI coordinate x=6, y=-70, z=44 (Kwok et al., 2012), and in a control area on the vertex, which was identified at the point of the same distance to the left and the right pre-auricular, and of the same distance to the nasion and the inion (Figure 1B). For combining each subject’s head with the MRI images, location information of each subject’s head was obtained individually by touching four fiducial points, which are the tip of the nose, the nasion, and the inter-tragal notch of each ear using an infrared pointer. The real-time locations of reflective markers which were attached to the coil and the subject were monitored by an infrared camera using a Polaris Optical Tracking System (Northern Digital, Waterloo, Canada).

In each session, TMS was delivered to either the precuneus or vertex before the main task. TMS was applied at 1 Hz frequency for a continuous duration of 20 min (1,200 pulses in total) at 110% of active motor threshold (MT), which was defined as the lowest TMS intensity delivered over the motor cortex necessary to elicit visible twitches of the right index finger in at least 5 out of 10 consecutive pulses (Rossini et al., 2015). The MT was measured both at the beginning of experiment session 1 in the memory and perceptual tasks. The order of stimulation sites was counterbalanced within subjects across tasks. During stimulation, participants wore earplugs to attenuate the sound of the stimulating coil discharge. The coil was held to the scalp of the participant with a custom coil holder and the subject’s head was propped a comfortable position. Coil orientation was parallel to the midline with the handle pointing downward. Immediately after the 20 min of rTMS, subjects performed four blocks of memory task in the MRI scanner (mean delay from rTMS to beginning of test: TMS-precuneus = 15.29 min, TMS-vertex= 20.76 min), or performed a visual perceptual task in a psychophysics room (mean delay from rTMS to beginning of test: TMS-precuneus = 6.7 min, TMS-vertex= 6.3 min). For safety reason and to avoid carry-over effects of rTMS across sessions, experimental sessions 1 and 2 were conducted on two separate days for both tasks (memory: mean interval = 8 days; perceptual: mean interval= 3.9 days).

## RESULTS

Overall, the participants missed 2.9% of TOJ trials and 2.2% confidence rating in the memory task, whereas the participants missed 0.7% trials in the perceptual type 1 task. Trials missing either one of the measures were excluded from the analysis.

We first examined whether the type 1 task performance in accuracy (% correct, Figure 1A), reaction time (RT, Figure 1B), and confidence rating (Figure 1C) might be affected by TMS. As expected, the task performance was not different between the two TMS conditions in neither memory (accuracy, *t* (17) = 0.349, *p* = 0.640; RT, *t* (17) = 1.997, *p* = 0.090; confidence rating, *t* (17) = 0.069, *p* = 0.780) nor perceptual part (accuracy, *t* (17) = 1.091, *p* = 0.480; RT, *t* (17) = 0.842, *p* = 0.490; confidence rating, *t* (17) = 0.461, *p* = 0.560).

We then used a robust metacognitive index (meta-d’ – d’) to investigate whether TMS on the precuneus might affect the metacognitive performance on the tasks. We performed a 2 (Task: memory/perception) × 2 (TMS: precuneus/vertex) repeated measures ANOVA for metacognitive efficiency – quantified as meta-d’ - d’ – from the SDT-based model and the hierarchical model separately. In the SDT-based model, we found an interaction effect between Task and TMS site (*F* (1, 17) = 7.25, *p* = 0.015; Figure 3A middle). The interaction was driven by lower metacognitive efficiency following TMS to precuneus relative to TMS to vertex in the memory task (*t* (17) = − 2.155, *p* = 0.046), whereas no difference in metacognitive efficiency was found in the perceptual task (*t* (17) = 1.378, *p* = 0.186). Metacognitive efficiency using the hierarchical model revealed the same pattern of results (Task × TMS interaction: *F* (1, 17) = 7.312, *p* = 0.015; memory: *t* (17) = −2.119, *p* = 0.049; perception: *t* (17) = 1.334, *p* = 0.200). To better characterize the effect of TMS on metacognitive efficiency, we performed sign tests to verify the extent of changes between TMS to precuneus and vertex. The metacognitive efficiency was reduced by TMS to precuneus in a majority of participants in the memory task (13/18 reduced, *p* = 0.035, sign test; Figure 3A left), but not in the perceptual task (10/18 reduced, *p* = 0.290, sign test; Figure 3A right).

These meta-indices are in principle based on how people rate their confidence, which refer to how meaningful a person’s confidence rating is in distinguishing between correct and incorrect responses. We accordingly ran a 3-way repeated measures ANOVA (Task: Memory/Perception × TMS: precuneus/vertex × Confidence: Low/High) on the type 1 task percentage correct and obtained a significant 3-way interaction (*F* (1, 17) = 10.652, *p* = 0.005). The TMS effect was disproportionally stronger in the memory task, as evident in a TMS × Confidence interaction (*F* (1, 17) = 4.487, *p* = 0.049; Figure 3B left), than in the perceptual task (*F* (1, 17) = 1.24, *p* = 0.281; Figure 3B right). Such effects in the memory task were driven by higher accuracy following TMS-precuneus than TMS-vertex in the low confidence ratings condition (*t* (17) = 2.354, *p* = 0.031), but not in the high confidence ratings condition (*t* (17) = −0.4, *p* = 0.694).

To add credibility to these results, we replicated these findings with the Phi coefficient (*F* (1, 17) = 13.81, *p* = 0.002; Figure 3C), confirming that our results were not biased by any idiosyncratic violations of the assumptions of SDT. These findings of lower metacognitive efficiency in the memory task following TMS to precuneus compared to vertex confirm our prediction that the precuneus causally mediates memory metacognition, but not perceptual metacognition.

To further probe whether the TMS effect on memory metacognition would be reflected by within-subjects changes in the between-tasks covariations, we calculated the between-tasks (Memory/Perception) correlations for all individuals’ type 1 task sensitivity (d’) and metacognitive efficiency respectively. We found that participants’ type 1 sensitivity (d’) between the perceptual and memory tasks are positively correlated, and that the magnitude of the correlation was not affected by TMS (TMS-vertex: *r* = 0.90, *p* < 0.001; TMS-precuneus: *r* = 0.82, *p* < 0.001; comparison between correlations: *z* = − 0.86, *p* = 0.390; Figure 4A). This again indicates that TMS had no effect on the basic task performance, in line with the pattern shown in Figure 2. In contrast, while the metacognitive efficiency for the two tasks were significantly correlated in the TMS-vertex condition (*r* = 0.72, *p* < 0.001; Figure 4B), as of what was reported previously (McCurdy et al., 2013), such correlational pattern was notably eliminated under TMS-precuneus treatment (*r* = −0.13, *p* = 0.63; Figure 4B), and the correlation coefficient was significantly lower than that of the TMS-vertex condition (*z* = − 3.38, *p* = < 0.001). Taken altogether, these results reveal that TMS to precuneus affects the metacognitive performance specifically for the memory domain.

**Figure 2:**
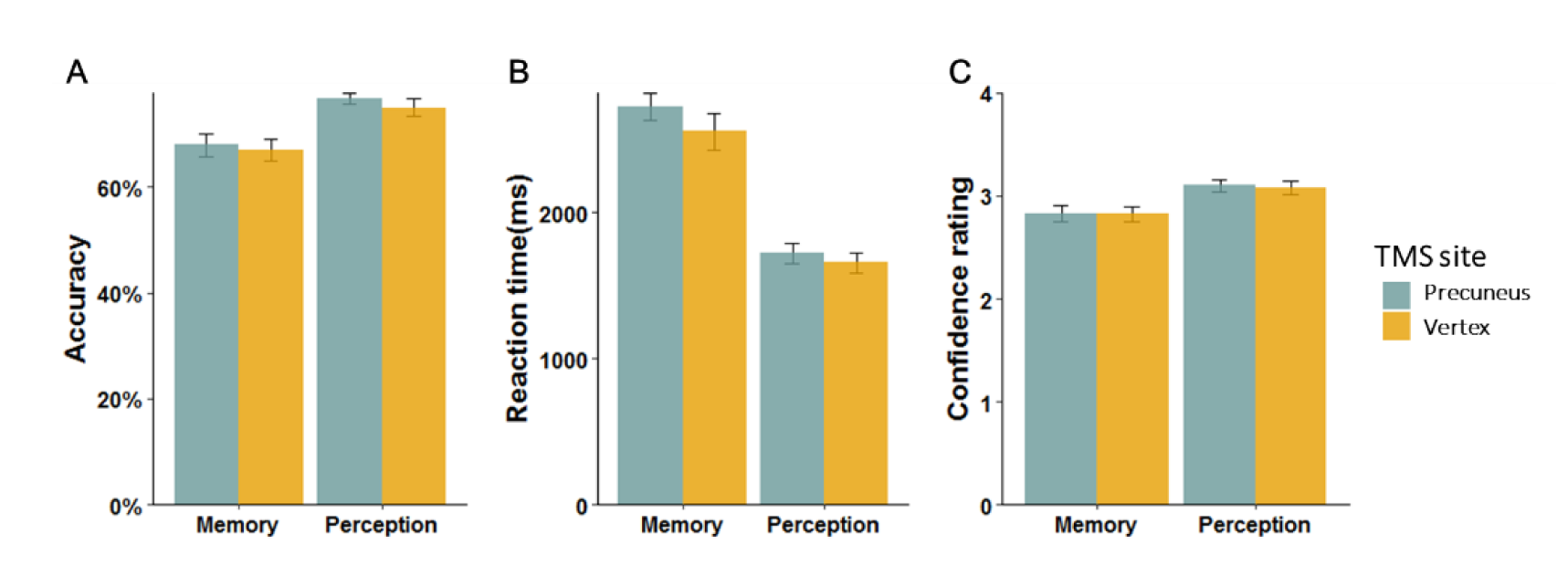
Basic task performance. Type 1 task performance was not affected by TMS in either of the tasks: **(A)** Accuracy **(B)** Reaction time **(C)** Mean level of confidence ratings. Error bars denote the standard error of the mean (SEM).

**Figure 3:**
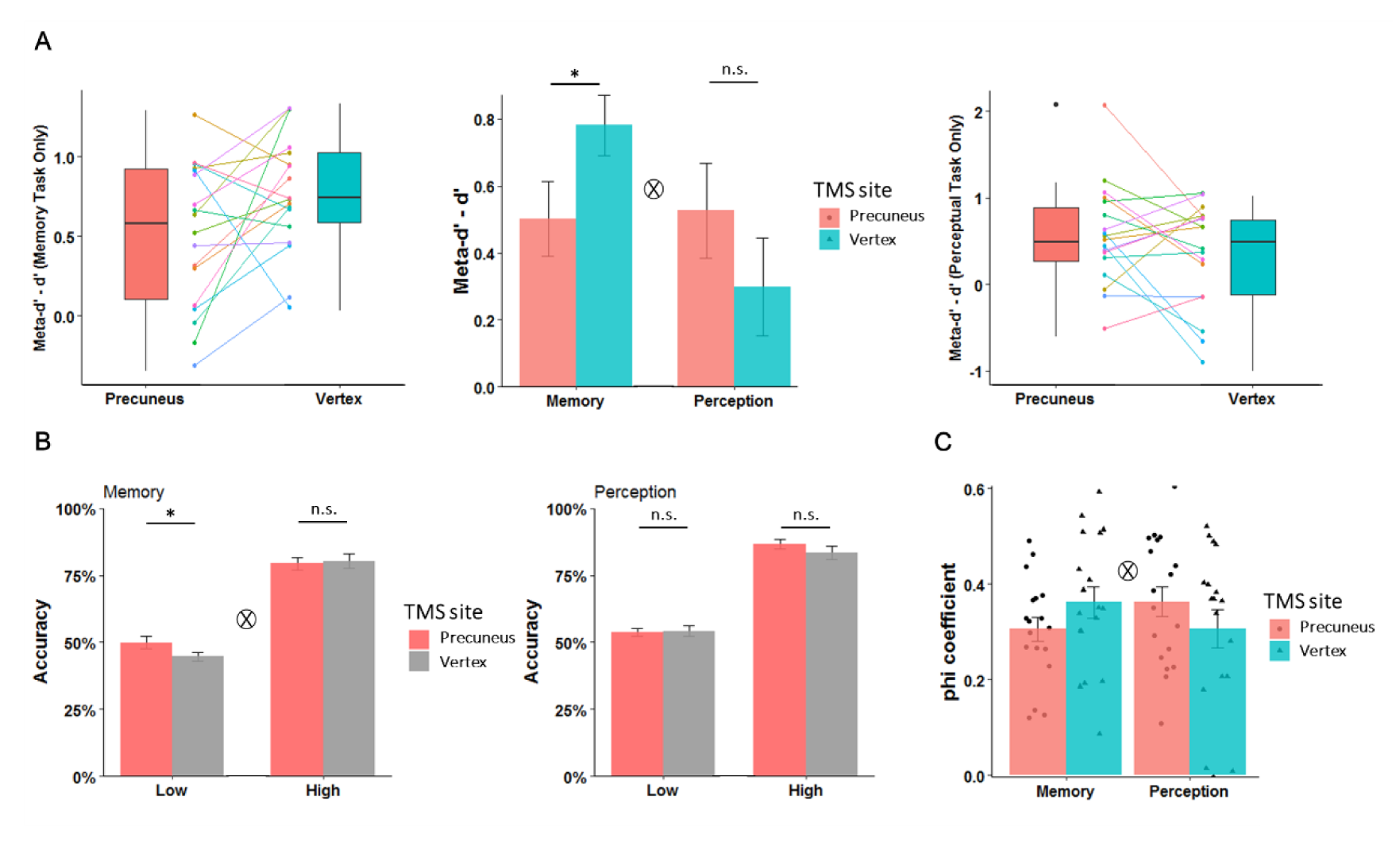
Differential effects of TMS on metacognitive performance. **(A)** Metacognitive efficiency in SDT-based model (meta-d’ - d’) under TMS-precuneus was lower than metacognitive efficiency under TMS-vertex in memory task but not in perceptual task. Each colored line depict within-subjects changes across conditions. **(B)** TMS ×Confidence ratings interaction in memory task. The accuracy for low confidence ratings under TMS-precuneus is significantly higher than that under TMS-vertex; no such effect for high confidence ratings. No significant effect of TMS in perceptual task. **(C)** TMS × Task interaction in phi coefficient. ⊗ indicates significant interaction *p* < 0.05, \**p* < 0.05, *n.s.* = not significant. Error bars represent SEM.

**Figure 4:**
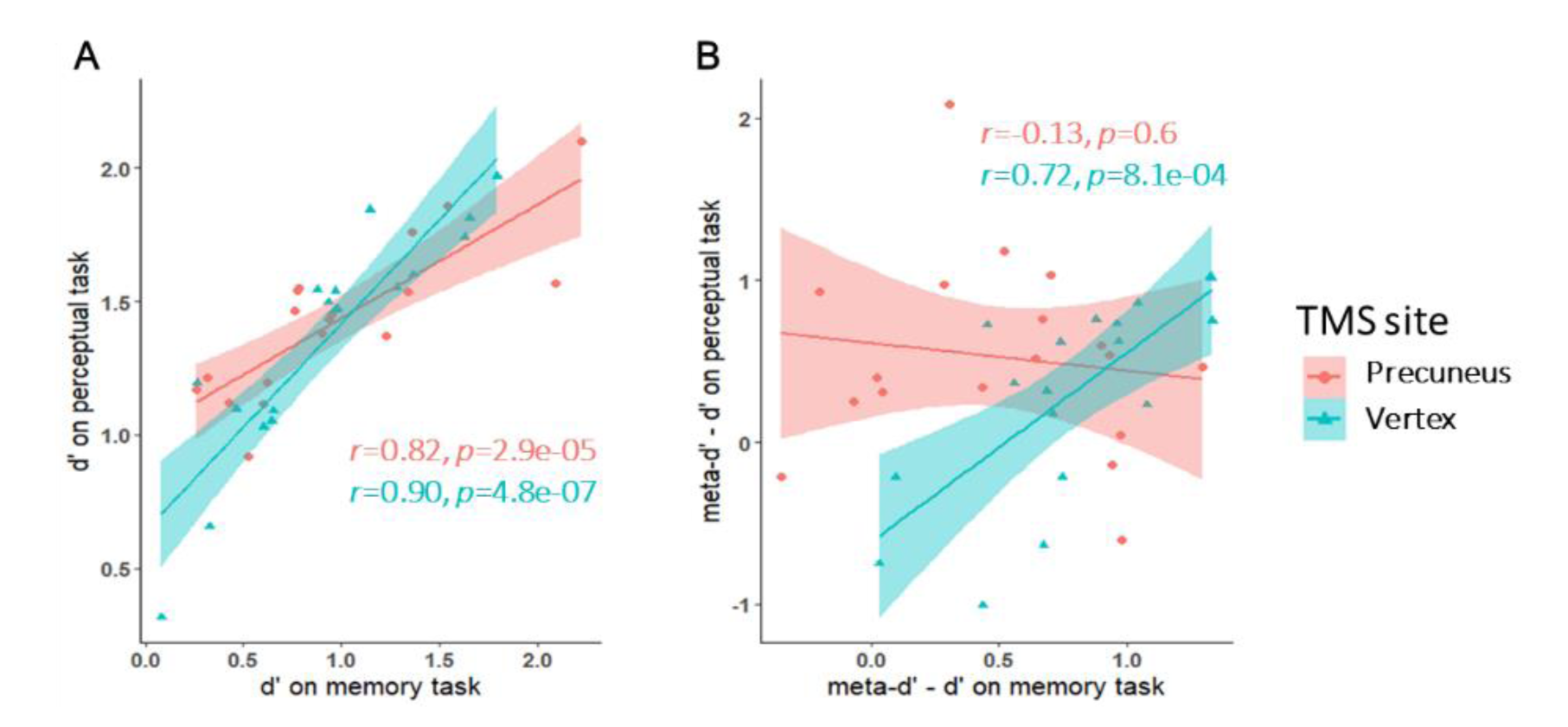
Correlation between memory and perceptual task performance and metacognitive indices. **(A)** TMS had no effect on the participants’ type 1 sensitivity (d’). The positive correlation between d’ on the perceptual and memory tasks was not affected by TMS. **(B)** In the TMS-vertex condition, the metacognitive efficiencies across the group were significantly correlated between memory and perceptual tasks. However, following TMS-precuneus, such between-tasks metacognitive efficiencies were no longer correlated.

## DISCUSSION

We employed an inferentially powerful technique to investigate the critical role of precuneus in the metacognitive ability in two distinct domains: memory and perception. We demonstrated that magnetic fields stimulation targeted at the precuneus impairs metacognitive efficiency in a long-term memory task without eliciting amnesia. TMS targeted to the precuneus affects the efficacy of confidence ratings specifically in a manner that subjects became less certain with their correct memory decisions. Critically, the TMS’s task-specific effect on the memory task, but not in the perceptual counterpart, implies that the neurobiological prerequisite for metacognitive ability is indeed supported by domain-specific components, some of which might be housed in the precuneus.

Previous studies showed that the precuneus is implicated in memory metacognition, derived from correlative measures such as anatomical connectivity and related functional activity analyses. For instance, a previous study identified a link between memory metacognitive efficiency and the precuneal gray matter density in healthy individuals (McCurdy et al., 2013), whereas a similar relationship was found between mnemonic metacognitive efficiency and functional connectivity between precuneus and medial aPFC (Baird et al., 2013). A recent study identified respective domain-specific and domain-general functional signals engaged by metacognitive judgments in perceptual and memory tasks using multivariate pattern analysis (Morales et al., 2018). They found that the domain-specific pattern for metacognition was encoded in the prefrontal cortex whereas the domain-general pattern was distributed in a widespread network in the frontal and posterior midline, including the precuneus. These studies thus suggest that the precuneus might be dually involved in both memory and perceptual metacognition for the close relationship shared between precuneus and perceptual metacognition (McCurdy et al., 2013; Morales et al., 2018). Considering that no prior study has executed controlled, targeted perturbation on this medial parietal region, we thus set out to examine its functional necessity for mnemonic metacognition by disrupting the precuneal function with TMS. Our TMS-induced focal disruption imposed a significant and selective effect on metacognitive ability in memory, but without altering the perceptual metacognitive performance at all. In a complementary manner, lesions to the anterior PFC are found to impair perceptual metacognitive ability while sparing the metacognitive efficiency for memory, indicating a domain-specific deficit in metacognition by anterior PFC lesions (Fleming et al., 2014). These findings conjointly provide causal evidence for a double dissociation in neural areas between memory and perceptual metacognition.

Our work carries implications for extending the metacognitive principle to episodic memory beyond the realm of working memory. The present finding is compatible with other human lesion and neuroimaging studies implicating the role of the parietal cortex in memory retrieval. For example, a lesion study showed that a patient with parietal cortex damage reporting that she felt less confident and experienced a lack of richness in the memories she retrieved (Davidson et al., 2008). This is consistent with other reports showing that lesions to the parietal cortex significantly diminish the retrospective confidence ratings, despite the performance remaining intact in a source recollection task (Simons, Peers, Mazuz, Berryhill, & Olson, 2010). Furthermore, in a functional neuroimaging study designed to tease apart different components of memory retrieval, activation in the precuneus was found to be associated with vividness judgments during episodic memory retrieval (Richter, Cooper, Bays, & Simons, 2016), consistent with the evidence that the precuneus serves to represent personally relevant content accompanied by vivid recollection (Sreekumar, Nielson, Smith, Dennis, & Sederberg, 2017) and detailed abstraction of temporal information required to support recollective TOJ (Ye et al., 2018). Given that vivid reminiscence is a defining feature of successful recollection of episodic events, the involvement of the precuneus during memory retrieval tasks might actually lie in its role in subserving the subjective experience of remembering. This argument aligns with the recent finding that EEG activity in the precuneus is linked with conscious dreaming experience (i.e., subjects remembered the content of dreaming experience after being awakened from a dream) (Siclari et al., 2017), in line with its role in mental imagery during retrieval (Fletcher et al., 1995). These behavioral and neural evidence convergently implicate the medial parietal cortex in the assessment of recollection during retrieval in support of its role in meta-memory. In line with the contribution to recollection of past episodes, our data corroborated the exiting evidence for the participation of precuneus in higher-order conscious processes during episodic memory retrieval.

Individual metacognitive efficiency scores were found to be positively correlated across the memory and perceptual domains under the control condition in some studies (Faivre et al., 2016; McCurdy et al., 2013; Ruby, Giles, & Lau, 2017; Samaha & Postle, 2017), but not in others (Baird et al., 2013; Vo et al., 2014; Fitzgerald et al., 2017; Sadeghi et al., 2017; Morales et al., 2018). It is plausible that such discord in correlation between metacognitive scores across domains is partly driven by the different types of judgments required (Ruby et al., 2017). A caveat is that the comparison did not take the stimulus characteristics across different tasks into account. Indeed, most studies of metacognition employed different categories of materials for the respective tasks, like word-list memory task versus dots-contained perceptual task (Baird et al., 2013; Fleming et al., 2014; McCurdy et al., 2013; Sadeghi et al., 2017). Following two recent studies (Ruby et al., 2017; Morales et al., 2018), here we also employed stimuli belonging to the same category – in fact identical sets of subject-specific stimuli material – for the memory and perceptual tasks, which would eliminate any confounds attributable to stimulus or featural characteristics.

To conclude, our findings reinforce the notion that precuneal region plays a critical role in mediating metacognition in episodic memory retrieval. To our knowledge, our study is the first one to causally verify the domain-specificity hypothesis of the precuneus in mnemonic metacognition in the human. Together with the contribution of anterior prefrontal cortex to perceptual metacognition, a challenge for future work is to understand how these different kinds of metacognition can be integrated into a unified framework.

## Acknowledgements

This work was supported by the Ministry of Education of PRC Humanities and Social Sciences Research grant 16YJC190006, STCSM Shanghai Pujiang Program 16PJ1402800, STCSM Natural Science Foundation of Shanghai 16ZR1410200, NYU Shanghai and the NYU-ECNU Institute of Brain and Cognitive Science at NYU Shanghai (S.C.K.). We thank Yudian Cai for his help in programming the perceptual task and Xinming Xu for suggesting resolution comparison for the perceptual task. Conflict of Interest: The authors declare no competing financial interests.

